# Pay to know me in your eyes: A computational account and oxytocin modulation of social evaluation

**DOI:** 10.1101/864330

**Authors:** Danyang Wang, Yina Ma

## Abstract

People are eager to know and recast the self in the eyes of others, even at a personal cost. However, it remains unknown what drives people to pursue costly evaluations of the self. Here, we propose that the evaluation of the self is valuable and that such subjective value placed on evaluation drives the costly-to-know behavior. By measuring the amount of money that individuals would forgo for the opportunity to know evaluations from other people (social evaluation) or a computer program (non-social evaluation), we quantified the subjective value individuals assigned to the evaluation on the self. The results from 5 studies (*n* = 375) lent cognitive and computational support for this hypothesis. Furthermore, the subjective value was modulated by the source and valence of the evaluation. Participants equally valued positive and negative non-social evaluations, characterized by a shared unknown aversion computation. However, individuals computed independent unknown aversion towards positive and negative social evaluations and placed a higher value on the opportunity to know another person’s evaluation on positive than negative aspects. Such a valence-dependent valuation of the social evaluation was facilitated by oxytocin, a neuropeptide linked to linked to social feedback learning and valuation processes, which decreased the value ascribed to negative social evaluation. Taken together, the current study reveals the psychological and computational processes underlying self-image formation and updating and suggests a role of oxytocin in modulating the value of social evaluation.

## Introduction

*He wants to see himself honestly, as others see him, but he also wants to see himself in the best possible light – Foundations of social psychology*

Humans are curious about how the self is evaluated ― either praised or criticized. Individuals spend over 70% of their daily conservation expressing their opinions about others and getting to know the self through the eyes of their friend, colleague, and employer^1–4^. We also frequently check social networking sites (such as Facebook and Twitter) to see how many people ‘like’ our pages or how others comment on our recent posts. However, it remains unknown what drives people to wonder how the self is evaluated, even at personal cost. In the current study, we propose that the subjective value individuals place on the opportunity to know evaluations of the self drives costly-to-know behavior. Three lines of research motivate this hypothesis. First, feedback on the self (e.g., praise and criticism) provides guidance for how to behave and/or interact appropriately with others^1,2^ and helps with individuals’ reputation formation and management^5,6^. Second, people do not tolerate ambiguity in self-related information; thus, the curiosity (of perceived self-image) provides an internal, psychological force driving individuals to explore how the self is evaluated^2,7,8^. Third, evaluative information conveys motivational value and has been used to motive expectation, learning, and decision-making^9,10^. Taken together, we hypothesized that high subjective value would be assigned to knowing evaluation.

To test this hypothesis, we develop a *pay-to-know* choice task where participants make a series of choices between two alternative options with different amounts of monetary reward that varied by the opportunity to know the evaluation or not. The *pay-to-know* choice task is modified from the “pay-per-view” paradigm. Using this paradigm, previous work has shown that viewing social signals or self-disclosure is intrinsically rewarding in that monkeys would sacrifice juice for the opportunity to view another monkey^11,12^; humans forgo monetary rewards to view attractive individuals of the opposite sex^13^ or to disclose information about oneself^14^. The current *pay-to-know* choice task allows us to measure the amount of money participants are willing to forgo to the opportunity to access evaluations from other people or a computer program and allows us to quantify the subjective value individuals assigned to knowing the evaluations. Moreover, integrating computational modeling, we sought to dissect the contribution of monetary reward and the opportunity to know evaluation in making choices in the *pay-to-know* task. Individual’s subjective cost of not knowing was characterized by a subject-specific unknown aversion parameter derived from a computational model of the individual’s choices.

Next, we asked the question of whether the subjective value assigned to the opportunity to know evaluation would be modulated by the valence of the evaluation aspects. The evaluation of the self functions to correct and guide one’s behavior^2,15^ and shapes one’s self-image^16^. Positive feedback reinforces the right behavior or choice, and negative feedback helps to avoid future error or failure; thus, both positive and negative evaluations of the self would be valuable. This could be especially true for evaluations based on objective standard criteria^17^. However, feedback has other functions besides telling right from wrong, especially social feedback. Social feedback (such as evaluation from other people) conveys different social signals depending on the valence. For example, positive social evaluation provides social support and social approval and facilitates social connection^18,19^ whereas negative social evaluation signals social threat and can hurt one’s feelings^20,21^. Thus, we hypothesize that the value of knowing non-social evaluation would be insensitive to valence, whereas people would be more motivated to know positive rather than negative social evaluations.

Finally, we aim to reveal the molecular substrates modulating the subjective value placed on evaluation. Oxytocin is of particular interest in the current study as oxytocin has been linked to social feedback learning and valuation processes^22,23^. Specifically, intranasal administration of oxytocin increased the learning of positive social feedback and decreased the learning of negative feedback. Moreover, animal work has shown that oxytocin decreased the value placed on the opportunity to view socially threatening stimuli (i.e., faces of dominant monkeys)^24^. These findings lead us to predict that individuals given oxytocin would value the positive social feedback to a greater degree and may devalue negative social feedback.

### Methods summary

#### The *pay-to-know choice task*

Participants were invited to trade off different amounts of monetary reward against the opportunity to have access to evaluations either provided by other people or a computer program (Fig. 1). Specifically, participants were first asked to provide one of their own photos and a self-introduction essay including their name, age, personality traits, likes/dislikes, hobbies, interests, etc. We then asked participants for permission to show their self-introduction to other people (strangers, Exp. 1, 3, 4) or enter it into a computer text/facial analysis program (Exp. 2, 3, 5) and told participants that other people or the computer program would make evaluations on them based on the self-introduction materials. Approximately 1 week later (range from 1 to 23 days), participants were then invited to the main experiment session and completed the *pay-to-know* choice task where participants chose between two options: ‘to-know’ (TK) and ‘not-to-know’ (NTK) the evaluations; these options differed in the amount of monetary reward received (payoff differences between ‘to-know’ and ‘not-to-know’ options, *Δm* = *M*_*TK*_ − *M*_*NTK*_, ranged from −3 to 3).

**Figure 1.**
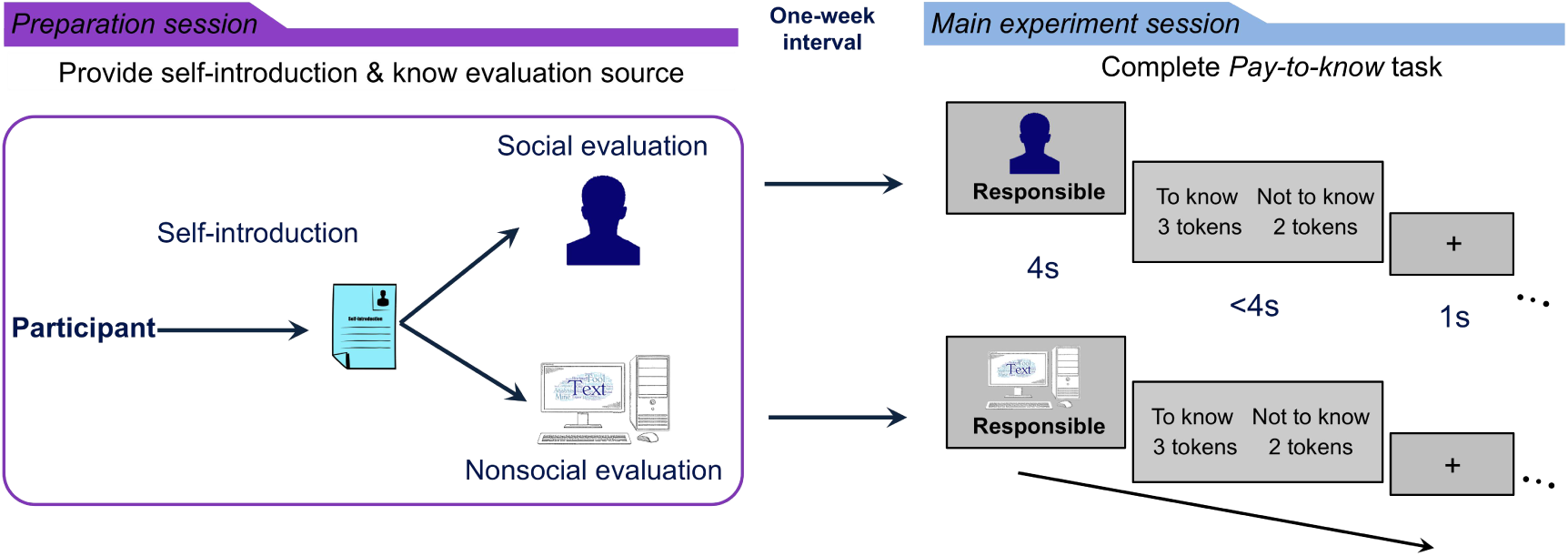
Illustration of the general experimental procedure. Participants were first invited to a preparation session to complete self-introduction as materials for social (i.e., other people) and non-social (i.e., a computer program) evaluations. Approximately 1 week later, participants came to the main experiment session and completed the *pay-to-know* choice task. For each trial of the *pay-to-know* task, participants were first presented with a single aspect (a positive or negative trait word) on which he could receive an evaluation and then had to decide whether they wanted to know the evaluation of the presented aspect or not. The ‘to-know’ (TK) and ‘not-to-know’ (NTK) options were associated with different monetary rewards (payoff difference ranged from −3 to 3).

#### Experimental design

We conducted 5 experiments (*n* = 375) in the current work. We first examined the subjective value individuals placed on the opportunity to know evaluations from other people (valuation of social evaluation, Exp. 1, *n* = 36) or from a computer program (valuation of non-social evaluation, Exp. 2, with an independent sample of *n* = 36). The comparison between Exp. 1 and 2 would reveal the shared and different valuations of social and non-social evaluations. In a third experiment, we sought to replicate the findings of the valuation of social and non-social evaluations observed in Exp. 1 and 2 in an independent online experiment with a larger sample (Exp. 3, *n* = 208, including the evaluation source (other people vs. computer program) as a between-subjects factor). We then employed a within-subject, double-blind, placebo-controlled design to examine the effect of oxytocin on the valuation of evaluations from other people (Exp. 4, *n* = 56) or computer programs (Exp. 5, *n* = 39). For all the experiments, we built computational models to fit participants’ choices in the *pay-to-know* choice task to reveal the computational processes that support the valuation of evaluation.

## Results

### Paying for the opportunity to know social evaluation

We first examined the extent to which participants would pay for the opportunity to know social evaluation (Exp. 1, *n* = 36). If knowing social evaluation is intrinsically motivating, we would expect participants to forgo monetary reward to choose the ‘to-know’ option; otherwise, participants would consistently choose whichever option was associated with higher monetary rewards to maximize their financial payoff. First, we found that participants generally chose to know social evaluations in more than 50% of the trials (overall knowing ratio: 63.57% ± 13.47% *vs*. 50%, *t*(35) = 6.05, *p* < 0.001, Fig. S1a). To exclude the possibility that the knowing ratio higher than 50% was driven by the choice of the ‘to-know’ option in the equal monetary amount condition (i.e., *M*_*TK*_ = *M*_*NTK*_), we calculated the knowing ratio for the trials where payoff amounts differed between the ‘to-know’ and ‘not-to-know’ options (referred to as the payoff-different knowing ratio). Among these payoff-different trials, participants still chose to know social evaluation in more than 50% of the trials (61.14 ± 13.66% *vs.* 50%, *t*(35) = 4.89, *p <* 0.001, Fig. S1b), confirming the value assigned to knowing social evaluation. For the trials where choosing the ‘to-know’ option gains no more money than choosing the ‘not-to-know’ option but is associated with monetary cost (i.e., *M*_*TK*_ ≤ *M*_*NTK*_, *costly-to-know* trials), participants chose costly to know social evaluations 43.33 ± 22.55% of the time (referred to as the costly knowing ratio). The overall knowing ratio (vs. 50%) and costly knowing ratio were henceforth reported to indicate whether and the extent to which individuals would prefer to know evaluations with a monetary cost. The analysis of the payoff-different knowing ratio showed the same pattern as the results of the overall knowing ratio (Fig. S1).

We then examined whether participants would equally prefer to know social evaluation on positive and negative aspects. If so, we would expect participants to forgo the same amounts of monetary reward to the opportunity to know either positive or negative aspects. We found that while participants valued the opportunity to know evaluations on both positive (69.15 ± 13.77% *vs.* 50%, *t*(35) = 8.35, *p <* 0.001) and negative (57.99 ± 19.11% *vs.* 50%, *t*(35) = 2.51, *p =* 0.017) aspects, participants preferred to know positive social evaluations more than negative ones (overall: *t*(35) = 3.42, *p =* 0.002, Fig. 2a; similar results of payoff-different knowing ratio, Fig. S1c). Moreover, participants more often chose costly-to-know positive (vs. negative) evaluations (higher costly knowing ratio: 49.01 ± 24.68% vs. 37.65 ± 26.42%, *t*(35) = 2.83, *p* = 0.008, Fig. 2b) and paid more money for the opportunity to know positive (vs. negative) evaluations (tokens cost on average: 0.65 ± 0.62 vs. 0.44 ± 0.51, *t*(35) = 3.27, *p* = 0.002, Fig. 2c).

**Figure 2.**
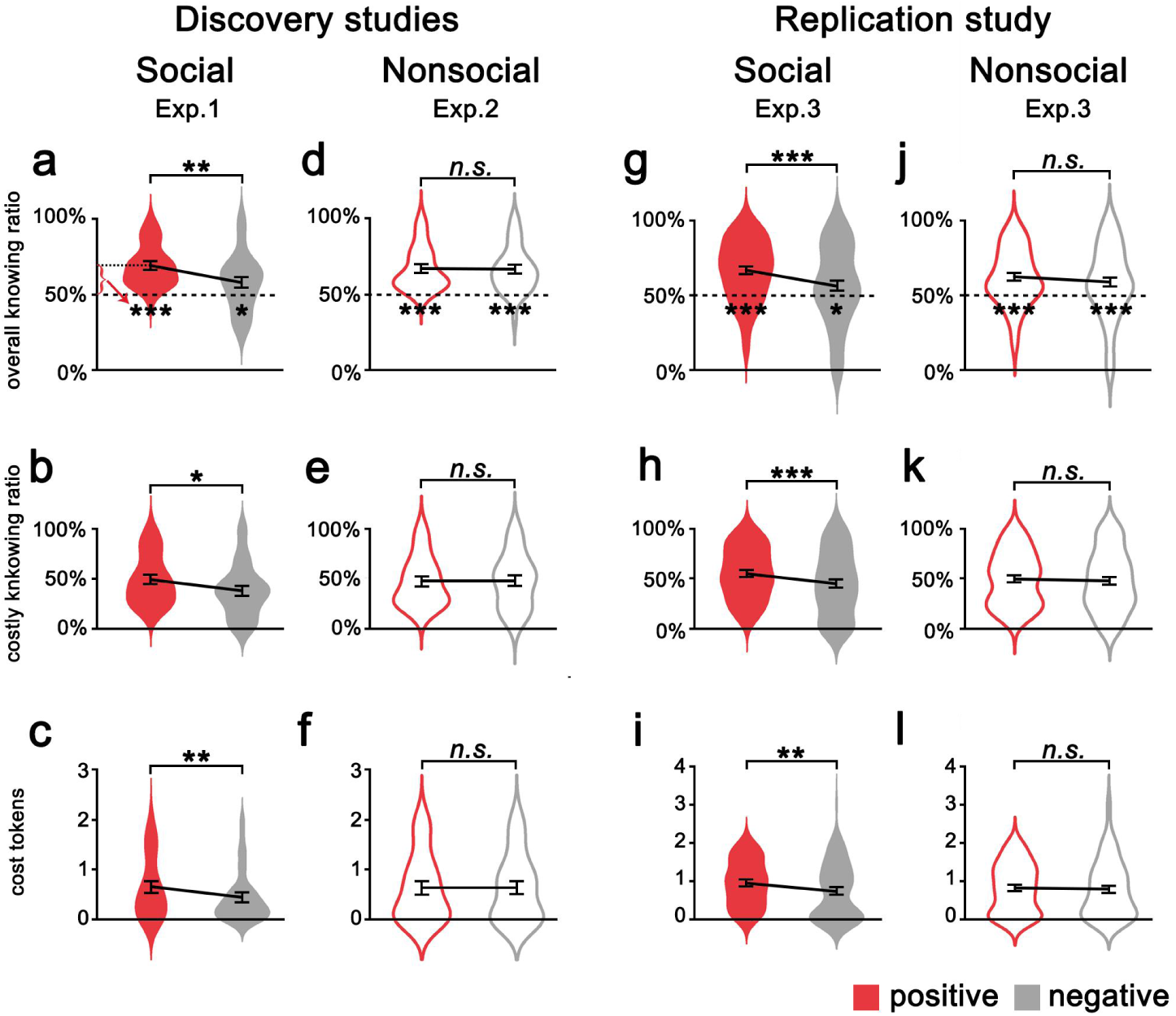
The model-free results in Exp. 1-3. Participants preferred to pay more to know social evaluations of positive than negative aspects (***a-c***) whereas they would forgo a similar amount of money to the opportunity to know positive and negative non-social evaluations (***d-f***). These results were replicated in online Exp. 3 (***g-l***). The violin plots indicate the distribution of indices from the *pay-to-know* choice task, with elements inside the violin plots representing the mean and standard error. (\**p* < 0.05, \*\**p* < 0.01 and \*\*\**p* < 0.001; *n.s.*, not significant)

### Paying to know social vs. non-social evaluations

Next, we asked whether participants placed subjective value on knowing social evaluation on themselves or on knowing any opinions of themselves. To address this question, we conducted Exp. 2 with an independent sample (Exp. 2, *n* = 36), following the same procedure as Exp. 1 except that participants were told that the evaluations they would receive were the outputs from the text/facial analysis of their self-introduction materials. We found that participants chose the ‘to-know’ option in 66.80 ± 13.49% of the trials (*vs.* 50%, *t*(35) = 7.48, *p* < 0.001, Fig. S1d, e). For the *costly-to-know* trials, participants chose costly ‘to-know’ 46.91 ± 24.21% of the time. Moreover, participants preferred to know the evaluations from the computer program on both the positive and negative aspects (66.97 ± 14.08% vs. 66.63 ± 15.16%, *t*(35) = 0.18, *p* = 0.859, Fig. 2d; similar results for payoff-different ratio, Fig. S1f). The cost of knowing evaluations on positive and negative aspects was comparable (costly knowing ratio: 46.48 ± 24.69% vs. 47.35 ± 27.39%, *t*(35) = −0.27, *p* = 0.790, Fig. 2e; tokens: 0.65 ± 0.67 vs. 0.63 ± 0.66, *t*(34) = 0.20, *p* = 0.842, Fig. 2f).

These results suggested that while individuals preferred to know evaluations both from other people and from a computer program, wanting to know social evaluation is valence-dependent whereas wanting to know non-social evaluation is valence-insensitive. To further confirm this difference, we conducted ANOVA on knowing ratio, with valence (positive vs. negative) as a within-subjects factor and the evaluation source (social vs. non-social) as a between-subject factor. This analysis showed a significant main effect of valence (*F*(1,70) = 9.30, *p* = 0.003) and a significant valence x source interaction (F(1,69) = 8.24, p = 0.005, Fig. 2a, d), confirming that the valence-dependent value was selective for social (but not non-social) evaluations. To specifically examine whether the willingness to know positive or negative evaluations was influenced by the evaluation source, we conducted simple effect analyses on the knowing ratio. We found that the knowing ratio for negative evaluation was lower for social sources than for non-social sources (*F*(1,70) = 4.52, *p* = 0.037), whereas the knowing ratio for positive evaluation was not influenced by the evaluation source (*F*(1,70) = 0.44, *p* = 0.508; Fig. S1c, f for similar results of payoff-different ratio). Similarly, 2 (valence: positive vs. negative) × 2 (source: social vs. non-social) ANOVAs on costly knowing behavior also revealed a significant main effect of valence (costly knowing ratio: *F*(1,70) = 4.14, *p* = 0.046; cost tokens: *F*(1,69) = 5.14, *p* = 0.027) and an interaction effect between valence and source (costly knowing ratio: *F*(1,70) = 5.63, *p* = 0.020, Fig. 2b,e; cost tokens: *F*(1,69) = 3.84, *p* = 0.054, Fig. 2c,f), as the effect of valence on costly behavior was similarly conditioned by the evaluation source. In addition, we asked participants to report the valence and self-relevance of each trait word, and there was no difference in the rating scores between the social and non-social evaluations (Exp. 1 vs. 2: *ps* > 0.10).

### Independent replication

Next, we performed an independent online experiment with a large sample (Exp. 3, *n* = 208), aiming to provide replication. Moreover, we additionally included a situation where both the ‘to-know’ and ‘not-to-know’ options were associated with monetary loss to test whether participants were not only willing to earn less but also to lose more money for the opportunity to know evaluation. We provided a replication of our findings that individuals indeed prefer costly ‘to-know’ evaluations from other people (in a valence-dependent way, to a greater degree for positive than negative aspects, Fig. 2g-i) and from a computer program (regardless of valence, equally for positive and negative aspects, Fig. 2j-l), both in the monetary gain and loss situations (Fig. S2). Given that no difference was found between the monetary gain and loss situations, we thus narrowed further analyses to monetary gain situation, and only included the monetary gain situation in Exp. 4 and 5.

### Quantifying the subjective value of evaluation

We next quantified the subjective value individuals placed on the opportunity to receive social and non-social evaluations by calculating the point of subjective equivalence (PSE) between ‘to-know’ and ‘not-to-know’ options. We plotted the proportion of ‘to-know’ choices against the monetary difference in the ‘to-know’ and ‘not-to-know’ options (i.e., *ΔM* = *M*_*TK*_ – *M*_*NTK*_, Fig. 3a, b) and fitted the data with sigmoid functions through a bootstrap procedure with 200 iterations (detailed in *Methods*) in the same way as the previous study^25^. The PSEs were derived from fitting sigmoid functions to each participant’s choices, representing the relative monetary value of ‘to-know’ over ‘not-to-know’ social (Fig. 3a) or non-social (Fig. 3b) evaluations. We showed that the PSE was significantly smaller than zero, suggesting that knowing social and non-social evaluations was worth significantly more than not knowing (Fig. 3c).

**Figure 3.**
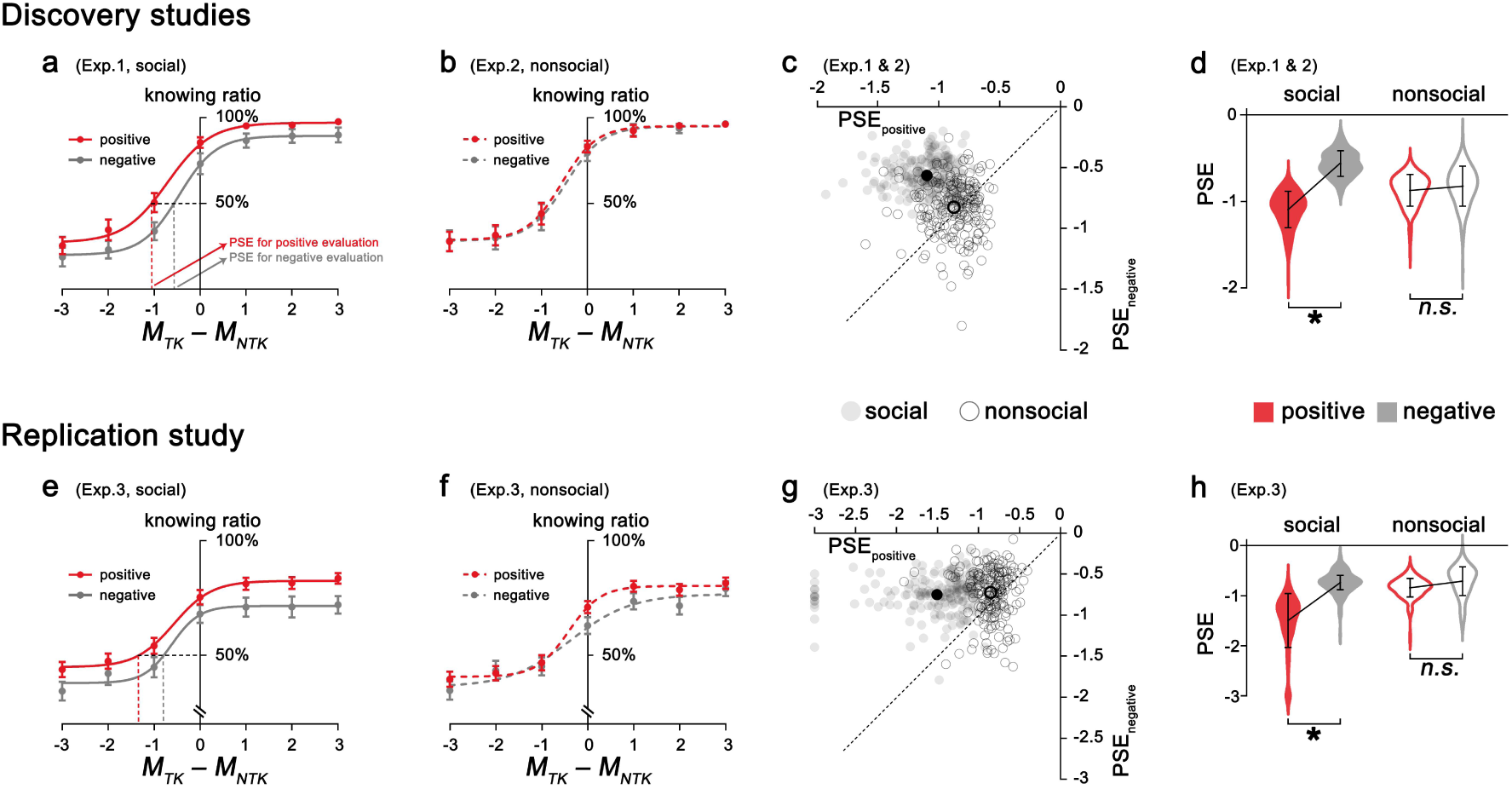
Subjective value assigned to social and non-social evaluations. The proportion of choosing to know the social (***a, e***) or nonsocial (***b, f***) evaluations was plotted against the monetary difference in the ‘to-know’ and ‘not to know’ options and fitted with sigmoid functions for the discovery and replication experiments, respectively (error bars represent standard errors across participants in each condition). The point of subjective equivalence (PSE) between ‘to-know’ and ‘not-to-know’ options was calculated. ***c*** (discovery sample) and ***g*** (replication sample) show the distributions of 200 bootstrapped sample means for social and non-social evaluations plotted against the PSE for evaluations of positive and negative aspects. Each gray dot represents the mean value of a bootstrapped sample, and dashed lines indicate the diagonal (the black dot indicates the mean value of 200 bootstrapped samples). Points on the diagonal represent bootstrapped samples with equal subjective value for knowing positive and negative evaluations. Points above the diagonal indicate bootstrapped samples with higher value assigned to knowing positive than negative evaluations, and points below represent samples assigning higher value to knowing negative than positive evaluations. The PSE of social evaluation was significantly higher for positive than for negative aspects, whereas the PSEs of non-social evaluation of positive and negative aspects were comparable (***d*** for discovery sample and ***h*** for the replication sample). The violin plots indicate the distribution of the PSEs, with elements inside representing the mean and bootstrap standard error (* and *n.s.* represent significant/insignificant in a bootstrap test).

The subjective value assigned to the opportunity to know evaluation equals earning 0.791 tokens on average for social evaluation (bootstrap 95% *confidence interval* (*CI*), −1.037 to −0.539) and 0.837 tokens for non-social evaluation (bootstrap 95% *CI*, −1.168 to −0.435). Moreover, the subjective value of social evaluation on positive aspects (PSE_positive_ = −1.057) was significantly higher than that for negative aspects (PSE_negative_ = −0.571, PSE_positive *vs*. negative_: bootstrap 95% *CI*, −0.861 to −0.101, Fig. 3c, d). However, the subjective values assigned to non-social evaluation for positive (PSE_positive_ = −0.838) and negative (PSE_negative_ = −0.811) aspects were comparable (PSE_positive *vs.* negative_: bootstrap 95% CI, −0.303 to 0.264, Fig. 3c, d). The same pattern was replicated in Exp. 3 (social: PSE_positive_ = −1.506 vs. PSE_negative_ = −0.749, bootstrap 95% CI, −0.966 to −0.114; non-social: PSE_positive_ = −0.809 vs. PSE_negative_ = −0.797, bootstrap 95% CI, −0.513 to 0.408; Fig. 3e-h).

### Computations underlying the valuation of evaluation

To reveal the computations underlying the *costly-to-know* behavior, we built a range of computational models to fit participants’ choices (*Supplementary Methods* for a list of related models), such as modified temporal-discounting models (which considered a temporal discount rate for the value of the evaluation^26–28^), and models that estimated loss aversion for monetary rewards. A softmax function was used to transform the trial-by-trial value into choice probability. We found that participants’ choices of the opportunity to know social or non-social evaluations were characterized by different computations.

Participants’ choices of knowing social evaluation or not were most parsimoniously explained by a model featuring distinct valuation of positive and negative social evaluations (i.e., *β*_*positive*_ and *β*_*negative*_), together with a parameter (i.e., *α*, ranging from 0 to 1) capturing the contribution of monetary payoff for the action choice (i.e., Model 5, Fig. 4a). Model 5, described in Fig. 4a, outperformed a series of alternative models (Table S1) and correctly predicted 88.25% of participants’ choices (95% *CI*, 85.64 to 90.86, Fig. 4a). The parameters *β*_*positive*_ and *β*_*negative*_ captured the impact of the desirability of knowing positive and negative evaluations on participants’ choices, respectively. The distinct valuation of knowing positive and negative social evaluations explained the model-free finding that participants traded different amounts of money to know social evaluation of positive and negative aspects.

**Figure 4.**
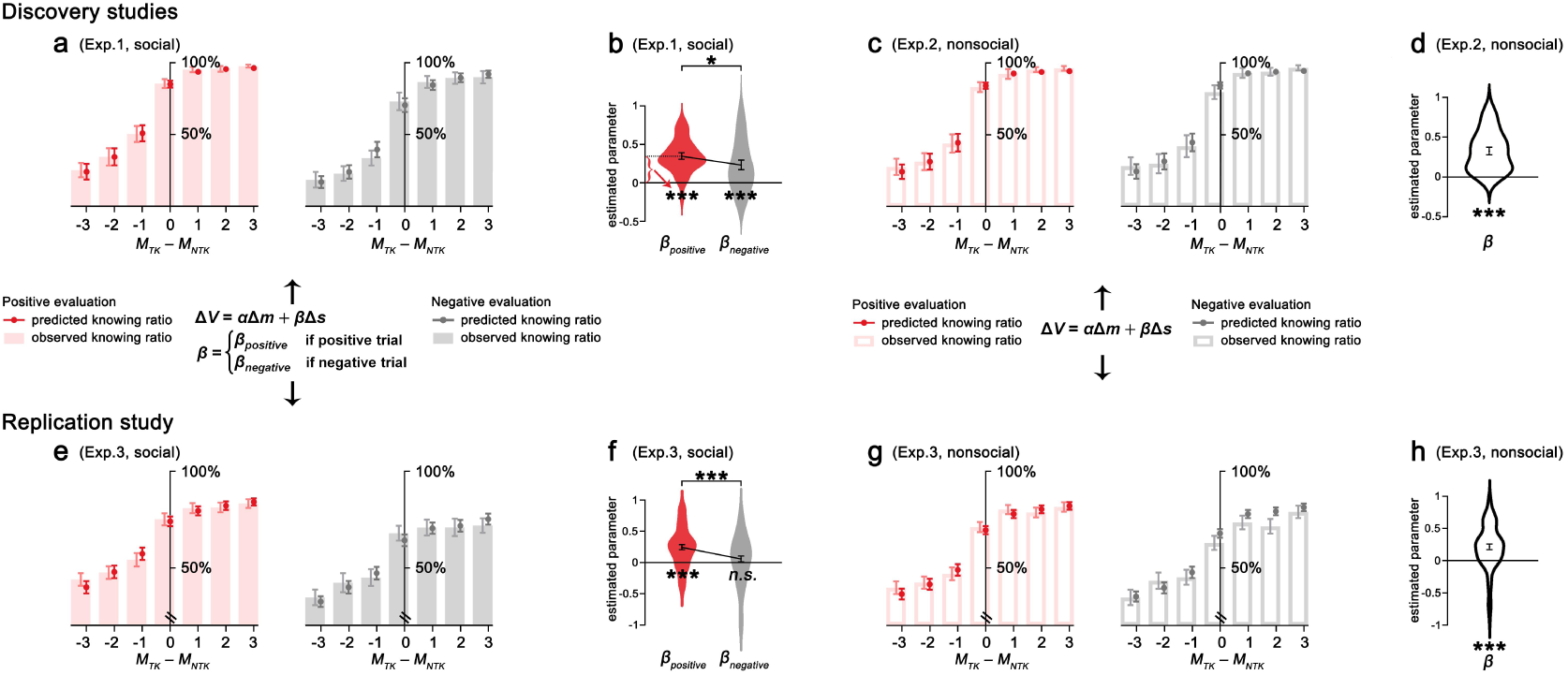
Computations underlying costly-to-know behavior. (***a, b***) Individuals’ choices in the pay-to-know social evaluation task were most parsimoniously explained by a model featuring the contribution of monetary payoff difference (*α*) and distinct contribution of knowing positive and negative social evaluations (*β*_*positive*_ > *β*_*negative*_) on the action choice. (***c, d***) In the non-social evaluation version, participants’ data were best fitted by a model featuring the valuation of monetary payoff difference (*α*) and unknown aversion towards non-social evaluation (*β* > 0). Observed data, circles; predicted choice of the ‘to-know’ option, bar charts (error bars represent standard errors across participants in each condition). The winning model and the related results were replicated in the independent replication experiment (**e-h**). The violin plots indicate the distribution of the estimated parameters, with elements inside representing the mean and standard error. (\**p* < 0.05, \*\*\**p* < 0.001, and *n.s.*, not significant).

It has been shown that while some individuals prefer to know the evaluations and avoid not knowing, others may show an aversion towards knowing evaluations, both for potentially positive^29^ and negative evaluations^30^. Thus, we set *β* parameters ranging from −1 to 1. Positive values of *β* indicated that participants were averse to not knowing (i.e., unknown aversion, *β* = 1 represents maximal aversion for unknowing, maximizing the value of the ‘to-know’ option), whereas negative values of *β* suggested that participants were unlikely to know the evaluation (*β* = −1 represents maximal aversion for knowing, minimizing the value of the ‘to-know’ option). The *β* parameter estimates derived from the winning Model 5 were significantly above 0 (*β*_*positive*_ = 0.35, *t*(35) = 9.57, *p* < 0.001; *β*_*negative*_ = 0.23, *t*(35) = 4.17, *p <* 0.001, Fig. 4b), suggesting unknown aversion for social evaluations separately on positive and negative aspects. Moreover, the *β* parameter estimates were higher for social evaluation of positive than negative aspects (*t*(35) = 2.31, *p=* 0.027, *d* = 0.38, 95% CI, 0.01 to 0.22, Fig. 4b). In addition, significantly more participants showed unknown aversion (i.e., *β* > 0) for social evaluations on positive (*N* = 35 out of 36) rather than negative (*N* = 26 out of 36) aspects (*χ*^*2*^ = 6.87, *p =* 0.009). These results suggested stronger aversion towards not knowing positive (relative to negative) social evaluations.

Participants’ responses in choosing whether to know the non-social evaluation or not were parsimoniously explained by Model 3 (outperformed Model 5), with two parameters accounting for the value of to-know non-social evaluation (*β*) and monetary reward (*α*). The winning Model 3 correctly predicted 87.75% of participants’ choices on whether to know non-social evaluation (95% *CI*, 84.04 to 91.45, Fig. 4c). The *β* parameter estimate derived from the winning Model 3 was significantly above 0 (*β* = 0.32, *t*(35) = 7.81, *p <* 0.001, Fig. 4d), suggesting unknown aversion for non-social evaluation. The shared unknown aversion for positive and negative aspects provided a computational explanation for the valence-insensitive subjective value of non-social evaluation. In addition, the winning model and the pattern of the parameter estimates derived from the winning model for both social and non-social evaluations were replicated in the independent online experiment (Exp. 3, *n* = 208, Fig. 4e-h, Table S1).

### The effect of oxytocin on the valuation of social evaluation

We then conducted Exp. 4 (*n* = 56, a within-subjects, double-blind, placebo-controlled design) to examine the modulation of intranasal oxytocin on the subjective value placed on the opportunity to know social evaluations. We first quantified the oxytocin effect on the subjective value placed on positive and negative social evaluations by comparing the PSEs under oxytocin and placebo. Oxytocin significantly decreased the subjective value of negative social evaluation (PSE_negative_, oxytocin vs. placebo: −0.326 vs. −0.625, bootstrap 95% *CI*, 0.033, 0.622, Fig. 5a, b), even to the extent of having no subjective value assigned to negative social evaluation, as the PSE was not significantly different from zero (PSE_negative-oxytocin_ = −0.339, bootstrap 95% *CI*, −0.626 to 0.040). However, the subjective value of positive social evaluation was not affected by oxytocin (PSE_positive_, oxytocin vs. placebo: −1.026 vs. −0.855, bootstrap 95% *CI*, −0.388 to 0.214, Fig. 5a, b).

**Figure 5.**
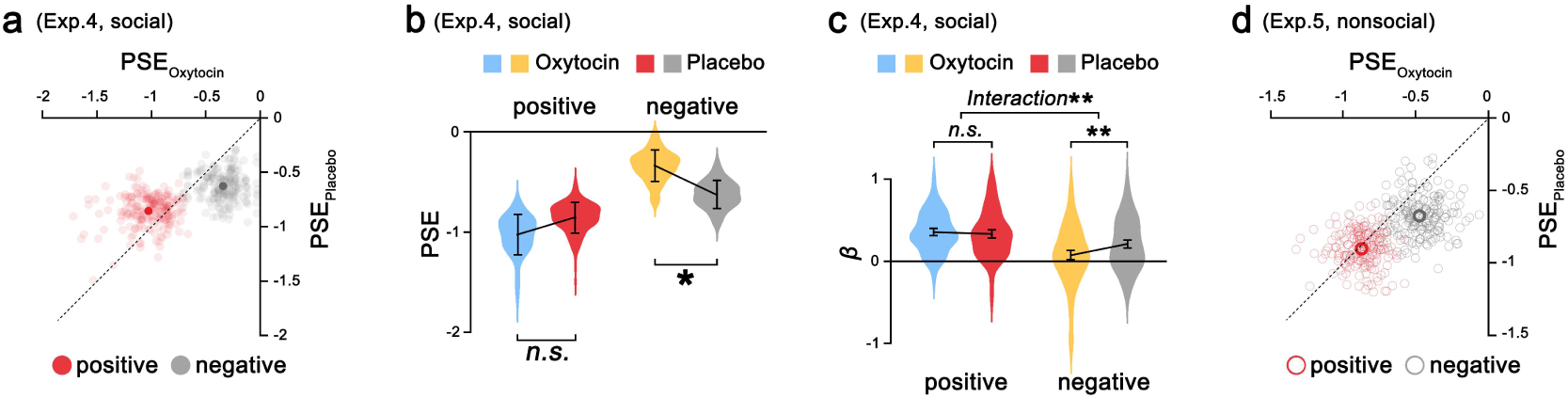
Oxytocin effect on the valuation of social evaluation. (***a, b***) The effect of oxytocin on subjective value placed on positive and negative social evaluations. Oxytocin decreased the point of subjective equivalence (PSE) for negative social evaluation but did not influence that of positive social evaluation. (***c***) Oxytocin decreased unknown aversion for negative social evaluation but not for positive social evaluation. (***d-f***) No significant effect of oxytocin was found for non-social evaluation. (\**p* < 0.05, \*\**p* < 0.01, and *n.s.* not significant). ***a*** (Exp. 4, social evaluation) and ***d*** (Exp. 5, social evaluation) show the distributions of 200 bootstrapped sample means for positive and negative evaluations plotted against the PSE in placebo and oxytocin treatment. Each dot represents the mean value of a bootstrapped sample, and dashed lines indicate the diagonal. Points on the diagonal represent bootstrapped samples placing equal subjective value on knowing evaluation in the oxytocin and placebo conditions. Points above the diagonal indicate bootstrapped samples with higher value assigned to knowing evaluation under oxytocin than placebo treatment, and points below the diagonal represent samples assigning lower value to knowing evaluation under oxytocin than placebo treatment. The violin plots indicate the distribution of the PSE, with elements inside representing the mean and bootstrap standard error (* and *n.s.* represent significant/insignificant in a bootstrap test).

Next, we examined the effect of oxytocin on the computational processes underlying the valuation of social evaluation. Participants’ choices regarding knowing social evaluation were also best fitted by Model 5 (Table S1, replicating Exp. 1 and 3). We found that intranasal oxytocin did not influence the weight of monetary payoff on participants’ choices (the *α* parameter estimates derived from Model 5, oxytocin vs. placebo: 0.33 vs. 0.35, *t*(55) = −0.89, *p=*0.379, Fig. S3). ANOVA on unknown aversion (*β*), with treatment (oxytocin vs. placebo) and valence (positive vs. negative) as within-subjects factors, showed a significant interaction (*F*(1,55) = 8.11, *p* = 0.006, Fig. 5c). While oxytocin did not significantly influence unknown aversion for positive social evaluation (*β*_*positive*_, oxytocin vs. placebo: 0.35 vs. 0.33, *t*(55) = 0.62, *p =* 0.541), oxytocin significantly decreased unknown aversion for negative social evaluation (*β*_*negative*_, oxytocin vs. placebo: 0.07 vs. 0.21, *t*(55) = −3.07, *p =* 0.003), even to the extent that participants given oxytocin did not show unknown aversion for negative social evaluation (*β*_*negative*_ under oxytocin was not significantly different from 0, *β*_*negative-oxytocin*_ = 0.07, *t*(55) = 1.45, *p* = 0.154).

To further examine whether a similar effect of oxytocin would be observed for non-social evaluation, we conducted Exp. 5 (*n* = 36, a within-subjects, double-blind, placebo-controlled design). Under placebo, we replicated the findings of Exp. 2 and 3 showing that participants valued non-social evaluation, and to a same extent for positive and negative aspects (Fig. 5d-f). Moreover, as predicted, no significant effect of oxytocin was observed on the valuation of non-social evaluation (Fig. 5d-f).

We also showed that mood changes before and after the experiment (Table S2, 3) and the decision times (Table S4) were not different between the oxytocin and placebo sessions in both Exp. 4 and 5. In addition, we asked participants to report the valence and self-relevance rating of each trait word, and no difference in rating scores was found between the oxytocin and placebo sessions (ps < 0.05). These results suggested that the effect of oxytocin on the valuation of social evaluation cannot simply be attributed to the potential effects of oxytocin on mood, cognitive performance or ratings of the evaluation aspects.

### Undervalued positive social evaluation in individuals with high depressive scores

We further examined individual differences in the valuation processes of social evaluation and in the effect of oxytocin on social evaluation. Specifically, we measured participants’ depressive scores by asking participants to complete the Beck Depression Inventory (BDI^31^), as depressive individuals have been shown to have a negative bias^32,33^ and reduced learning of positive social feedback^34^.

We first examined the relationship between healthy individual’s depressive score and the valuation processing of social evaluation by correlating participants’ BDI scores and unknown aversion under placebo. We found that under placebo, the BDI scores were negatively correlated with unknown aversion for positive social evaluation (indicated by *β*_*positive*_ derived from social evaluation model, *r* = −0.351, *p* = 0.008, Fig. 6a) but not for negative social evaluation (*r* = −0.15, *p* = 0.282, Fig. 6b), suggesting less unknown aversion for social evaluation (positive ones in particular) in individuals scored higher in depressive symptoms. Interestingly, this relationship was dampened by oxytocin. Unknown aversion did not vary significantly with BDI scores after oxytocin administration (*ps* > 0.3). Furthermore, the oxytocin effect on *β*_*positive*_ (i.e., *β*_*positive-oxytocin*_ – *β*_*positive-placebo*_) was positively correlated with depression scores (*r* = 0.269, *p* = 0.045, Fig. 6c), suggesting stronger oxytocin effects on increasing aversion towards not-knowing positive social evaluation in more depressive individuals. In addition, the BDI score was not related to unknown aversion for non-social evaluation under placebo (*r* = −0.061, *p* = 0.710), nor did it modulate the oxytocin effect (*r* = 0.019, *p* = 0.910).

**Figure 6.**
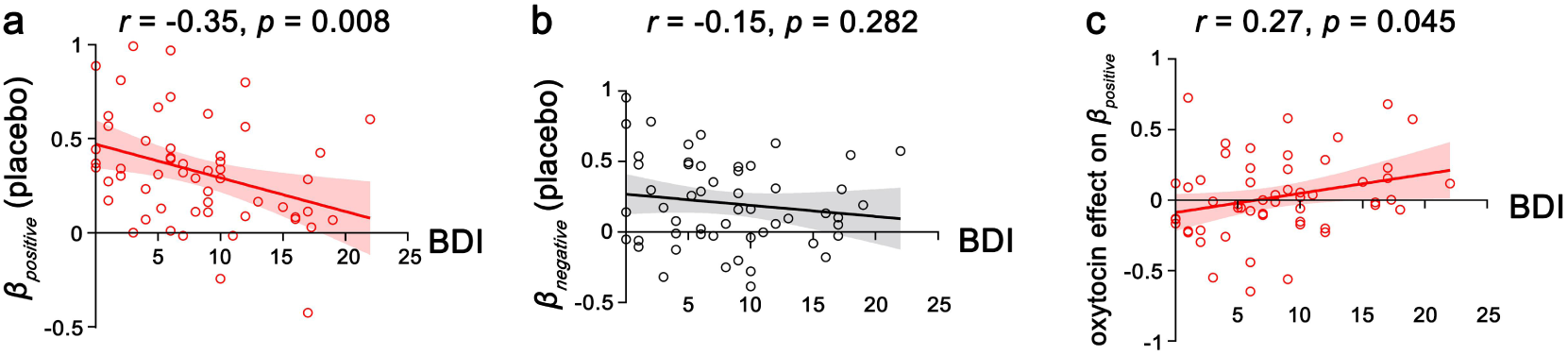
Individual differences in the valuation of social evaluation and the related oxytocin effect. (***a*, *b***) Under placebo, individuals scored high on the Beck Depression Inventory (BDI) showed less unknown aversion for positive (***a***) but not negative (***b***) social evaluations. (***c***) Stronger effects of oxytocin on increasing the valuation of positive social evaluation were found in individuals who scored higher on the BDI. Each solid line represents the least squares fit with shading showing the 95% CI.

## Discussion

Knowing how the self is perceived and evaluated helps individuals to form and update self-image, appropriately interact with others, and fit in the social world^2,15,16,18,19^. The current study examined whether and how individuals value the opportunity to know opinions on the self. We demonstrate the subjective values individuals placed on knowing the evaluation of the self and further reveal how such subjective value is computed and modulated by the evaluation source (i.e., evaluation from other people or a computer program) and valence (i.e., evaluation on positive or negative aspects of the self). We show that whether the subjective value of evaluation is modulated by valence depends on the evaluation source. Specifically, the value assigned to positive and negative non-social evaluations is comparable and computed via a shared unknown aversion. In contrast, the value placed on positive and negative social evaluations is computed via independent unknown aversion parameters, resulting in higher value assigned to knowing positive social evaluation. Taken together, we show evidence that people care about the image of the self in others’ eyes to the extent that they would forgo monetary reward to the opportunity to know evaluations of the self.

The willingness to forgo monetary reward to the opportunity to know indicated individual’s intrinsic motive to know evaluation of the self. The next question is what drives such a motive. To quantify the driving force behind the costly-to-know behavior, we built a range of computational models and identified an individual-specific unknown aversion parameter (although varied widely but exhibited in most individuals), as a key determinant of the willingness to know despite monetary cost. Receiving positive evaluations is associated with self-approval or perceived as social reward^1,35,36^, whereas receiving negative evaluation could guide individual’s future behavior. Thus, being unknown may serve as a psychological cost associated with losing the opportunity of knowing the self, as being unknown about the evaluation may cause uncertainty about how others view the self and how to behave in future social interactions or to manage one’s reputation^37^. It should be noted that although it has been shown that individuals are motivated to receive positive feedback and that being praised is rewarding^1,35,36^, our finding of placing subjective value on the opportunity to know evaluation of the self could not be simply explained by the motivation related to being praised. First, all evaluations were designed to be given after the experiment rather than immediately after participants chose the ‘to-know’ option. Second, in the current study, individuals were willing to forgo monetary reward to know evaluations of both positive and negative aspects.

Moreover, the driving force behind wanting to know social and non-social evaluations might be different. Individuals’ choices for the opportunity to know non-social evaluation were characterized by the unknown aversion computation shared by positive and negative aspects, i.e., participants’ choices of and the subjective value assigned to non-social evaluation were independent of the valence of the evaluation aspects. When facing non-social evaluation, individuals are motivated to acquire an accurate self-image by knowing how the self is objectively evaluated, as we would expect objective evaluations from a computer program. However, wanting to know social evaluation may not be driven by seeing an objective self in others’ eyes. Individuals computed the unknown aversion separately for positive and negative social evaluations and placed lower value on the negative ones. While positive social feedback conveys social approval and conformity^19,38^, negative social feedback signals social rejection^20,21^. Thus, being unknown about negative social feedback would help us avoid potential social rejection or hurt.

The evaluation of the self is an important way to be aware of how the self is perceived and to build self-image^17,34,39^. An individual’s self-image is formed and updated as a result of how the individual sees the self, as well as how others see the individual, which are respectively linked to the private and public self-image^40,41^. Public self-image, reflecting the self in the presence of others^42^, is mainly formed and updated based on social evaluations. Our finding of the positively biased valuation of social evaluation (i.e., higher subjective value and stronger unknown aversion for positive social evaluations) may provide a cognitive path to facilitate a positive public self-image. This is further supported by the result of undervalued positive social evaluation in individuals with high depressive scores, who are less motivated to manage a positive public self-image^43,44^. On the other hand, findings of equal subjective value and shared unknown aversion for positive and negative non-social evaluations implied the need for an objective and unbiased self-image in private^17,45^. Thus, our finding furthered our understanding of public and private self-images in that public and private self-images are distinguished by the different goals of managing a positive self-image in public and maintaining an accurate self-image in private.

Interestingly, we observed an oxytocin effect on the valuation of social (rather than non-social) evaluation, decreasing the subjective value and unknown aversion for negative social evaluation to the extent of having no subjective value and no unknown aversion towards negative social evaluation. This was consistent with previous animal and human studies of the attenuated processing of negative social information by oxytocin. For example, oxytocin dampened learning of negative social feedback^22^ and the influence of negative facial expressions on behaviors^24,46^. Although oxytocin did not influence the valuation of positive social evaluation in general, we observed a modulatory effect of depressive score on the oxytocin effect, showing a selective effect of oxytocin on increasing unknown aversion of positive social evaluation in individuals with higher depression scores. These findings may implicate the potential of oxytocin in alleviating social anhedonia and blunted responses to social reward, typical symptoms of depression^47–51^. This is also consistent with previous findings of a stronger oxytocin effect in less socially adapted individuals^52,53^.

## Methods

### Ethics Approval

The experimental procedure was in line with the standards set by the Declaration of Helsinki and was approved by a local ethics committee at the State Key Laboratory of Cognitive Neuroscience and Learning, Beijing Normal University (Beijing, China). All participants provided written informed content after the experimental procedures had been fully explained and acknowledged their right to withdraw at any time during the experiment. The current study was pre-registered via the Open Science Framework with all materials and data available at https://osf.io/f6djw/.

### Participants

We recruited 375 paid males in the current study. All participants were healthy, right-handed, free of a history of neurological or psychiatric disorders, and had normal or corrected-to-normal vision. Given that previous studies reported sex differences in oxytocin effects on social cognition and behavior^53,54,55^, we only recruited male participants for the current study. In addition, in the discovery experiments (Exp. 1 and 2) and the oxytocin experiments (Exp. 4 and 5), we only recruited single males to avoid the potential influence of romantic relationship status on social motivation^56^. In the replication experiment (Exp. 3), we recruited participants regardless of their romantic relationship status and further showed that *costly-to-know* behavior was not modulated by romantic relationship status (Table S5, 6).

### Sample size estimation

We conducted sample size estimation using G*Power 3.1.9.2^57^ to calculate the number of participants needed for the experiments in the laboratory (Exp. 1, 2, 4, and 5) to detect a reliable effect with α = 0.05 and power = 0.8. Given that no previous studies have examined the valuation of choosing to know evaluations, we based on a medium effect size of *Cohen’s d* = 0.5^58,59^ to estimate the sample size for Exp. 1 and 2 (which were conducted in parallel) and revealed that a sample of 34 participants was needed to detect a reliable effect when testing the knowing ratio against that occurring by random choice (50%) in a one-sample t-test. Exp. 1 and 2 each recruited a sample of 36 participants (Exp. 1, age = 22.36 ± 2.75 years; Exp. 2, age = 22.72 ± 2.39 years). In the online experiment (Exp. 3), as an online replication of our results, we recruited a large sample of 208 participants (age = 22.24 ± 3.49 years).

For the oxytocin experiment (Exp. 4), the sample size estimation was based on an estimated effect size revealed in a meta-analysis of oxytocin effects^60^ (*Cohen’s d* = 0.35). The G*Power calculation suggested that 52 participants were needed to detect a reliable interaction effect in a 2 (valence: positive vs. negative)-by-2 (treatment: oxytocin vs. placebo) within-subjects design. We thus recruited 56 participants (age = 21.21 ± 2.76 years) in Exp. 4. Based on the effect size from the original finding in Exp. 4 (*Cohen’s d* = 0.41), we conducted power analysis and predetermined that 37 participants were required for a reliable effect. Thus, 39 participants were recruited in Exp. 5 (age = 21.82 ± 2.95 years).

### General procedure

In Exp. 1-3, participants were invited to two separate sessions (with a mean interval of 1 week, Fig. 1). In the first session, participants provided self-introduction materials, based on which evaluation would be made. In the second session, participants completed mood-related questionnaires, the *pay-to-know* choice task, and a post-rating survey.

For the oxytocin experiments (Exp. 4 and 5), we employed a double-blind, placebo-controlled, within-subjects design where participants come to the oxytocin and placebo sessions (at least 7-day interval, mean interval = 10.9 ± 6.3 days, with treatment order counterbalanced across participants) after the first self-introduction session. For the oxytocin and placebo sessions, participants were instructed to refrain from smoking or drinking (except water) for 2 hours before the experiment. Upon arrival, participants first completed the Positive and Negative Affect Schedule^61^ to measure pre-treatment mood. Approximately 35 min before the main *pay-to-know* choice task, participants intranasally self-administered a single dose of 24 IU oxytocin or placebo (containing the active ingredients except for the neuropeptide) under an experimenter’s supervision. The procedure of oxytocin and placebo administration was similar to that used in the previous work that showed reliable effects of oxytocin on social cognition and behaviors^22,23,62^. The spray was administered to participants three times, with each administration consisting of one inhalation of 4 IU into each nostril. A 24 IU dose of oxytocin, commonly used in the oxytocin literature^23,63^, was employed in the current study because it has been recently shown to produce more pronounced effects (than 12 or 48 IU) on socio-affective processing, as well as on plasma and saliva oxytocin levels^64^.

In the *pay-to-know* choice task, participants completed two 63-trial blocks (with 63 positive traits and 63 negative traits, matched on arousal rating). For each trial (Fig. 1), participants were first presented with a trait word and the evaluation source (the face of a stranger or the text-analysis program) for 4 s. Participants were then asked to choose between two options within 4 s, i.e., a ‘to-know’ option and a ‘not-to-know’ option that were associated with different amounts of tokens.

At the end of the second session of Exp. 1-3 and the end of each oxytocin or placebo session, participants completed a post-experiment rating of the trait words (including valence and self-relevance rating).

### Data analyses

In each experiment, we asked participants to rate the self-relevance of all 126 trait words (−10 = not like me at all, 10 = exactly like me). On the basis of the self-relevance rating, we divided the trait words into self-relevant and self-irrelevant traits for each participant. To prevent spurious confounds of self-relevance on the valuation process^3^, formal data analysis was performed on the self-relevant trait words on a participant basis.

For the model-free indices, we mainly focused on the overall knowing ratio, payoff-different knowing ratio, costly knowing ratio, and cost tokens. All the indices were calculated separately for trials of positive and negative traits. 1) The overall knowing ratio was the proportion of choosing the ‘to-know’ option in all trials; 2) the payoff different knowing ratio was the proportion of choosing the ‘to-know’ option in trials where the payoff differed between two options; 3) the costly knowing ratio was the proportion of choosing the ‘to-know’ option in costly-to-know trials where choosing the ‘to-know’ option gains no more money than choosing the ‘not-to-know’ option; 4) the cost token was calculated as the averaged amount of tokens associated with the unchosen ‘not-to-know’ option in costly-to-know trials.

To quantify the subjective value participants placed on knowing, each participant’s choice data was fit with sigmoid functions to give an estimation of the point of subjective equivalence (PSE) between the ‘to-know’ and ‘not-to-know’ options.

The sigmoid function is defined as follows:

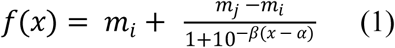

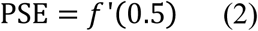

where *x* represents the payoff differences between ‘to-know’ and ‘not-to-know’ options (i.e., *∆M* = *M*_*TK*_ – *M*_*NTK*_), *f(x)* represents the knowing ratio in each payoff difference condition, *m*_*i*_ represents the lower asymptote, *m*_*j*_ represents the upper asymptote, *β* represents the slope of the sigmoidal response function, and *α* represents the x-value of the sigmoid midpoint. The PSE was calculated as the monetary value at which a participant effectively chose arbitrarily between the ‘to-know’ and ‘not-to-know’ options (equation 2). The estimated PSEs were obtained through a bootstrap procedure with 200 iterations, similar to the previous work^25^. Sigmoid curves were fit to the knowing ratio for each bootstrap sample by implementing nonlinear least-squares estimates in R. The goodness of fit of each bootstrap sample to the sigmoid function was measured, and the sigmoid function fitted well with each sample (r^2^ ranging from 0.932 to 1.000 across all experiments).

### Computational Modeling

We fit a range of models to each participant’s choice data to capture the valuation process of choosing to know evaluation on a trial-by-trial basis. For each model, trial-by-trial value differences (i.e., *∆V*) were transformed into choice probabilities using a softmax function^65,66^:

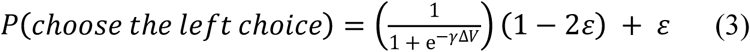

where γ is an inverse temperature parameter capturing the sensitivity of choices to *ΔV* and *ε* is a lapse rate that captures choice noisiness resulting from factors independent of *ΔV*. These parameters were estimated for each participant by implementing nonlinear optimization in MATLAB (MathWorks, Inc.).

A Bayesian model comparison technique^67,68^ was employed to compare models. For each model, we computed the Bayesian Information Criterion (BIC) scores for each participant. We then compared the group BIC scores (summed BIC scores across participants) and identified the best-fit model with the lowest group BIC score. We showed that Model 5 (3), for social (non-social) evaluation choices, outperformed a range of alternative models, including those considering temporal discounting across trials or monetary loss aversion (Table S1).

## Supplementary Information

### Supplementary Methods

#### Trait word selection

We first conducted a pilot study to select trait words for the main experiment. We aimed to identify positive and negative trait words that were different in valence but matched on arousal. Thus, we recruited 33 male participants (age: 22.88 ± 3.35) and asked them to rate on the valence (i.e., how negative or positive a trait word is) and arousal (i.e., the intensity of a trait word) of 307 trait words that were selected on the basis of a comprehensive list of trait adjectives^1^ and the Chinese Personality Adjective List^2^. The valence and arousal were rated on a 9-point Likert scale (from −4 to 4, for valence: extremely negative to extremely positive; for arousal: very calming to highly exciting). Based on the rating scores, we included 63 positive (e.g., responsible, handsome, romantic, knowledgeable) and 63 negative words (e.g., lazy, greedy, immature, superficial) in the main experiments. The positive and negative trait words were significantly different in valence (t = 15.38, p < 0.001) but matched on arousal ratings (t = 1.53, p = 0.136).

### Data analysis

#### List of alternative models for model comparison

To arbitrate the computational processes employed by the participants, we compared a range of models, each of which explained choices in terms of the value difference (*ΔV*) between the left and right choices. Models 1 through 6 differed in model complexity, mainly capturing the contribution of monetary payoff difference and the contribution of knowing the evaluation on action choice. More complex model variants included independent contributions of monetary payoff differences and to-know evaluation, and separate parameters for the positive and negative trials. In Models 7 and 8, we assumed that the participant’s choices changed over the course of the session and considered a parameter that captured temporal discounting of the subjective value difference between the ‘to-know’ and ‘not-to-know’ options. In Models 9 and 10, we conceptualized a loss aversion towards monetary reward, assuming that participants require more money for choosing not-to-know than they are willing to pay to know.

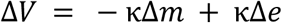

##### Model 1

In Model 1, the likelihood of choosing the left choice is a function of the value difference (*ΔV*) between the two choices. The value difference depends on the difference in monetary payoff (*Δm* = *M*_*left*_ – *M*_*right*_) and to-know evaluation or not (*Δe* = 1, if left choice is ‘to know’; *Δe* = −1, if left choice is ‘not to know’) and an unknown aversion parameter that captures the subjective cost of not-knowing evaluation, which is independent of the valence of the evaluation. When *κ* approaches 1, participants are maximally averse to not-knowing; as *κ* approaches −1, participants are maximally averse to knowing evaluation.

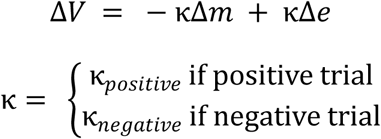

##### Model 2

In Model 2, we assumed that participants made decisions by separately evaluating the costs of not-knowing evaluations for positive and negative aspects, considering independent unknown aversion parameters for positive and negative aspects (i.e., *κ*_*positive*_ and *κ*_*negative*_, respectively).

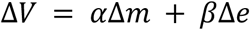

##### Model 3

Model 3 tested whether participants make decisions based on separate evaluation of the contribution of the monetary payoff differences and to-know evaluations or not to action choice. In this model, we obtained separate estimates for *α* and *β*.

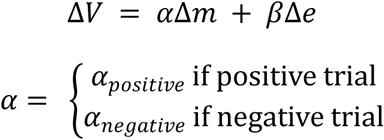

##### Model 4

Model 4 was similar to Model 3 in that it allowed for the separated contribution of monetary payoff difference and knowing evaluations but further tested whether participants considered the contribution of monetary payoff to different degrees when choosing for positive and negative aspects.

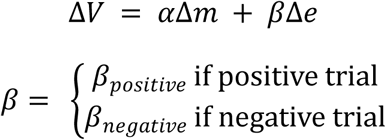

##### Model 5

Model 5 tested whether participants were averse to not-knowing evaluations of positive and negative aspects to a different degree. In this model, we obtained separate estimates for *β*_*positive*_ and *β*_*negative*_.

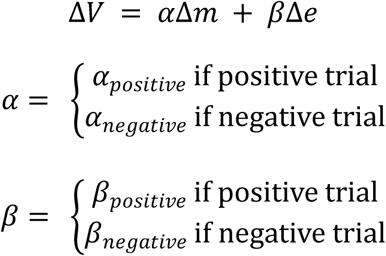

##### Model 6

Model 6 was similar to Model 5 but further tested whether participants considered different degrees of the contribution of monetary payoff differences on action choice for positive and negative trials.

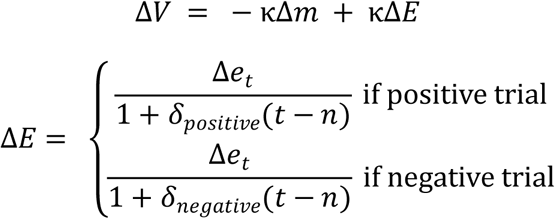

##### Model 7

Model 7 tested the possibility that subjective value differences between the ‘to-know’ and ‘not-to-know’ options would be discounted over the course of the session by considering the temporal discounting of the subjective value difference. It is possible that participant’s motivation for choosing to know evaluation is decreased due to the fatigue effect. In model 7, *Δe*_*t*_ on trial *t* was hyperbolically discounted at a discount rate *δ*; *n* is the total number of trials. We also tested whether subjective value differences between to-know and not-to-know positive and negative evaluations would be independently discounted by considering independent discount rates for positive and negative evaluations.

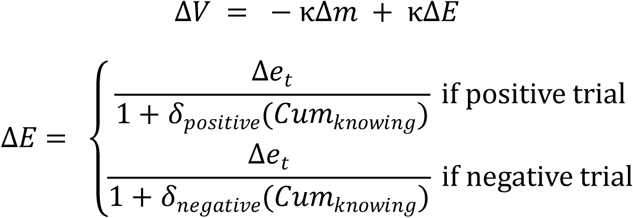

##### Model 8

Model 8 was similar to Model 7, but this model assumed that the change in a participant’s preference would be dependent upon whether more ‘to-know’ choices were made rather than more ‘not-to-know’ choices were made, as in Model 7. It is possible that the participant’s preference towards knowing evaluations is weakened over the course of the session because the satisfaction increased after making enough ‘to-know’ choices. In model 8, *Δe*_*t*_ on trial *t* was hyperbolically discounted at a discount rate that was independent of positive and negative aspects, *δ*_*positive*_ and *δ*_*negative*_; *Cum*_*knowing*_ represented the accumulative frequency of ‘to-know’ choices.

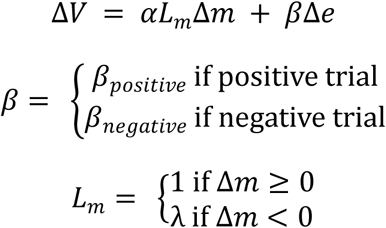

##### Model 9

Model 9 was similar to Model 5 but further tested whether participants were loss-averse for monetary payoff (*λ*). Note that loss aversion, in the context of the current experiment, produces a pattern of choices in which participants require more money to forgo knowing evaluations than they are willing to pay to know when the ‘to know’ option is associated with larger monetary payoff, and participants require more money to choose knowing evaluations than they are willing to pay for not-to-know when the ‘to-know’ option is associated with smaller monetary payoff. This effect is consistent with an omission bias in moral decision making^4^.

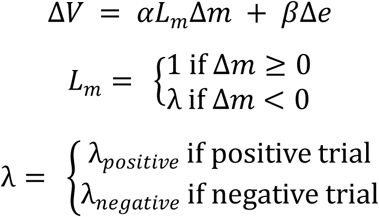

##### Model 10

Model 10 is similar to Model 3 but further tested whether participants showed different degrees of loss aversion for monetary payoff in positive and negative trials.

---

**Table S1.** Bayesian Information Criterion (BIC) for all models.

| Models | Parameters | No. Of<br>Parameter<br>s per<br>participan<br>t | Model<br>BIC<br>Exp1 | Model<br>BIC<br>Exp2 | Model<br>BIC<br>Exp3<br>social | Model<br>BIC<br>Exp3-nons<br>ocial | Model<br>BIC<br>Exp4<br>PL | Model<br>BIC<br>Exp5<br>PL |
| --- | --- | --- | --- | --- | --- | --- | --- | --- |
| Model 1 (M1) | $\kappa$ | 3 | 3689.24 | 3164.42 | 9508.37 | 10122.87 | 5564.71 | 3524.01 |
| Model 2 (M2) | $\kappa_{positive}, \kappa_{negative}$ | 4 | 3724.52 | 3224.94 | 9633.11 | 10249.43 | 5588.09 | 3580.82 |
| Model 3 (M3) | $\alpha, \beta$ | 4 | 2205.95 | **<u>1944.32</u>** | 7031.25 | **<u>7979.17</u>** | 3439.63 | **<u>2144.55</u>** |
| Model 4 (M4) | $\alpha_{positive}, \alpha_{negative}, \beta$ | 5 | 2241.54 | 1986.44 | 7123.71 | 8090.72 | 3483.68 | 2193.48 |
| Model 5 (M5) | $\alpha, \beta_{positive}, \beta_{negative}$ | 5 | **<u>2173.03</u>** | 1967.87 | **<u>6974.80</u>** | 7985.59 | **<u>3367.77</u>** | 2147.13 |
| Model 6 (M6) | $\alpha_{positive}, \alpha_{negative}, \beta_{positive}, \beta_{negative}$ | 6 | 2234.15 | 2028.86 | 7118.95 | 8166.79 | 3468.92 | 2214.99 |
| Model 7 (M7) | $\kappa, \delta_{positive}, \delta_{negative}$ | 5 | 2918.96 | 2579.80 | NaN | NaN | 4321.65 | 2800.74 |
| Model 8 (M8) | $\kappa, \delta_{positive}, \delta_{negative}$ | 5 | 2909.65 | 2737.54 | NaN | NaN | 4363.82 | 3053.44 |
| Model 9 (M9) | $\alpha, \beta_{positive}, \beta_{negative}, \lambda_m$ | 6 | 2249.56 | 2049.69 | 7146.16 | 8145.09 | 3493.08 | 2235.12 |
| Model 10 (M10) | $\alpha, \beta, \lambda_{m-positive}, \lambda_{m-negative}$ | 6 | 2281.62 | 2112.86 | 7224.42 | 8220.10 | 3595.47 | 2258.58 |

Note:
Model 5 (3) fitted best participants' choices in the social (non-social) pay-to-know task in a model comparison that considers differences in model complexity. More complex model variants included separate parameters for the positive-trait and negative-trait conditions, discount rate for outcome, and loss aversion for monetary payoff. The Bayesian Information Criterion (BIC) scores are the Bayesian equivalent to a fixed effects analysis.

**Table S2.**
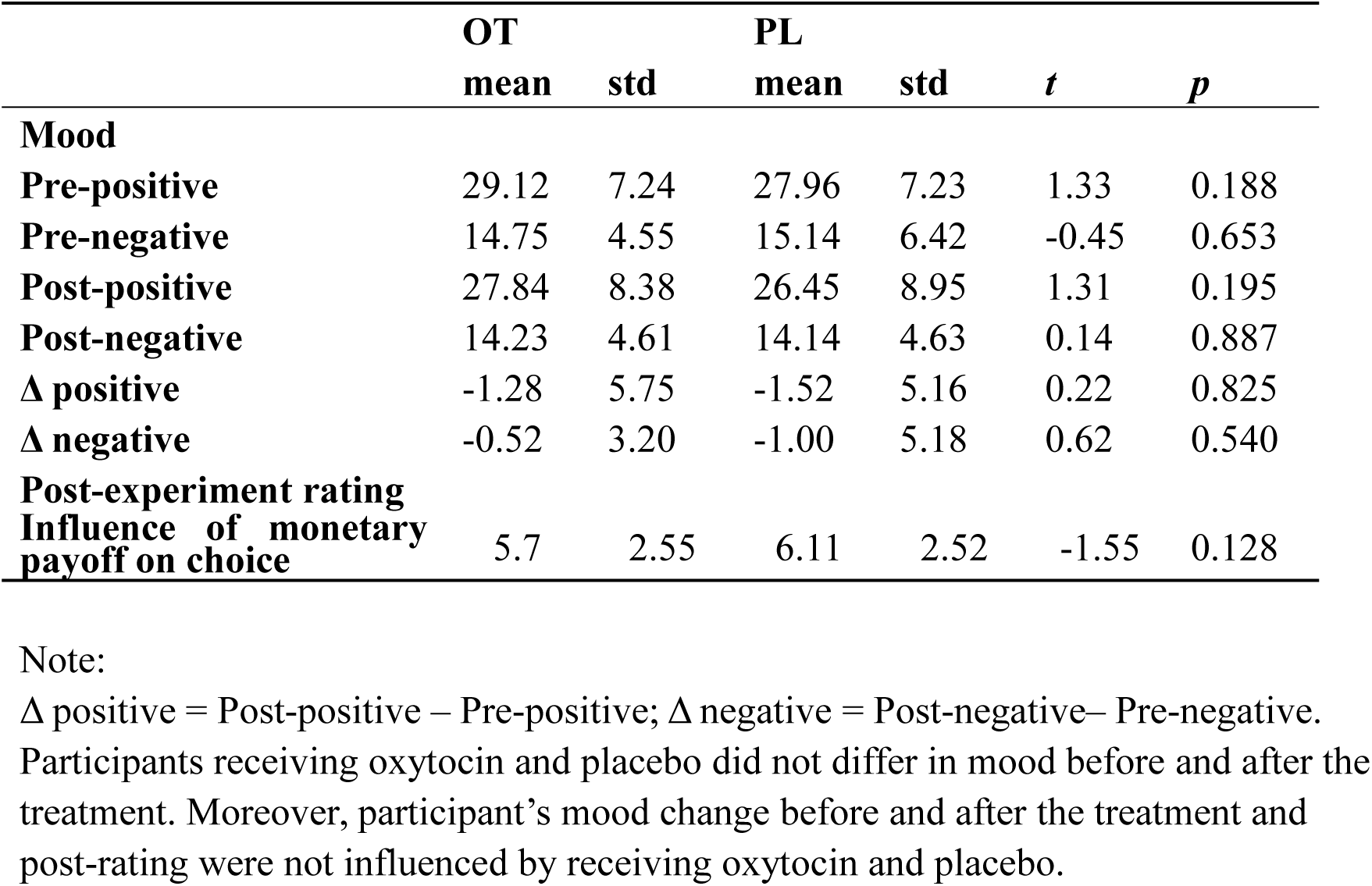
Pre-experiment and post-experiment mood, mood change, and post-experiment rating scores on attitude under oxytocin and placebo administration in Exp. 4

**Table S3.** Pre-experiment and post-experiment mood, mood change, and post-experiment rating scores under oxytocin and placebo administration in Exp. 5

|  | OT |  | PL |  |  |  |
| --- | --- | --- | --- | --- | --- | --- |
|  | mean | std | mean | std | *t* | *p* |
| **Mood** |  |  |  |  |  |  |
| **Pre-positive** | 31.61 | 6.18 | 33.11 | 5.50 | -1.72 | 0.093 |
| **Pre-negative** | 14.66 | 5.26 | 14.53 | 5.76 | 0.18 | 0.862 |
| **Post-positive** | 28.56 | 7.88 | 30.69 | 7.30 | -1.93 | 0.062 |
| **Post-negative** | 15.89 | 5.59 | 15.11 | 6.30 | 0.95 | 0.351 |
| **Δ positive** | 3.28 | 5.64 | 2.53 | 5.20 | 0.76 | 0.451 |
| **Δ negative** | -1.72 | 4.15 | -1.31 | 4.57 | -0.38 | 0.705 |
| **Post rating** |  |  |  |  |  |  |
| **Influence of monetary payoff on choice** | 6.15 | 2.39 | 6.15 | 2.19 | <0.01 | 1 |

Note: Δ positive = Post-positive – Pre-positive; Δ negative = Post-negative– Pre-negative. Participants receiving oxytocin and placebo did not differ in mood before and after the treatment. Moreover, participant’s mood change before and after the treatment and post-rating were not influenced by receiving oxytocin and placebo.

**Table S4.** Mean reaction times (RTs, ms) under oxytocin and placebo administration in Exp. 4 and 5

| Experiment | Groups | All trials | Positive trials | Negative trials |
| --- | --- | --- | --- | --- |
| **Exp. 4** | PL: mean (std) | 1120.22<br>(341.43) | 1119.69<br>(341.34) | 1120.76<br>(352.36) |
|  | OT: mean (std) | 1110.81<br>(337.15) | 1109.30<br>(322.35) | 1112.33<br>(632.18) |
| | PL vs. OT: $t$<br>($p$) | 0.26 (0.796) | 0.29 (0.770) | 0.21 (0.837) |
| **Exp. 5** | PL: mean (std) | 1354.51<br>(500.28) | 1349.90<br>(474.49) | 1359.13<br>(541.08) |
|  | OT: mean (std) | 1370.82<br>(396.26) | 1360.73<br>(392.63) | 1380.91<br>(442.66) |
| | PL vs. OT: $t$<br>($p$) | -0.31 (0.757) | -0.19 (0.848) | -0.37 (0.714) |

**Table S5.** Modulation of romantic relationship status on decision-making for social evaluation in Exp.3

| Variables | *F* | *p* |
| --- | --- | --- |
| Overall knowing ratio (positive trials) | 0.75 | 0.526 |
| Payoff-different knowing ratio (positive trials) | 0.78 | 0.506 |
| Costly knowing ratio (positive trials) | 1.38 | 0.253 |
| Cost tokens (positive trials) | 1.39 | 0.250 |
| Overall knowing ratio (negative trials) | 0.48 | 0.696 |
| Payoff-different knowing ratio (negative trials) | 0.32 | 0.811 |
| Costly knowing ratio (negative trials) | 1.19 | 0.317 |
| Cost tokens (negative trials) | 0.83 | 0.482 |
| Model-based indices: $\alpha$ | 2.85 | 0.041 |
| Model-based indices: $\beta_{positive}$ | 0.19 | 0.900 |
| Model-based indices: $\beta_{negative}$ | 1.69 | 0.175 |

Note: The *F* and *p* values were from one-way ANOVA

**Table S6.** Modulation of romantic relationship status on decision-making for non-social evaluation in Exp.3

| Variables | *F* | *p* |
| --- | --- | --- |
| **Overall knowing ratio (positive trials)** | 0.94 | 0.424 |
| **Payoff-different knowing ratio (positive trials)** | 0.70 | 0.553 |
| **Costly knowing ratio (positive trials)** | 0.44 | 0.727 |
| **Cost tokens (positive trials)** | 0.54 | 0.657 |
| **Overall knowing ratio (negative trials)** | 0.12 | 0.950 |
| **Payoff-different knowing ratio (negative trials)** | 0.28 | 0.839 |
| **Costly knowing ratio (negative trials)** | 0.79 | 0.504 |
| **Cost tokens (negative trials)** | 1.30 | 0.280 |
| **Model-based indices: $\alpha$** | 0.50 | 0.681 |
| **Model-based indices: $\beta$** | 0.69 | 0.562 |

Note: The *F* and *p* values were from one-way ANOVA

**Figure S1.**
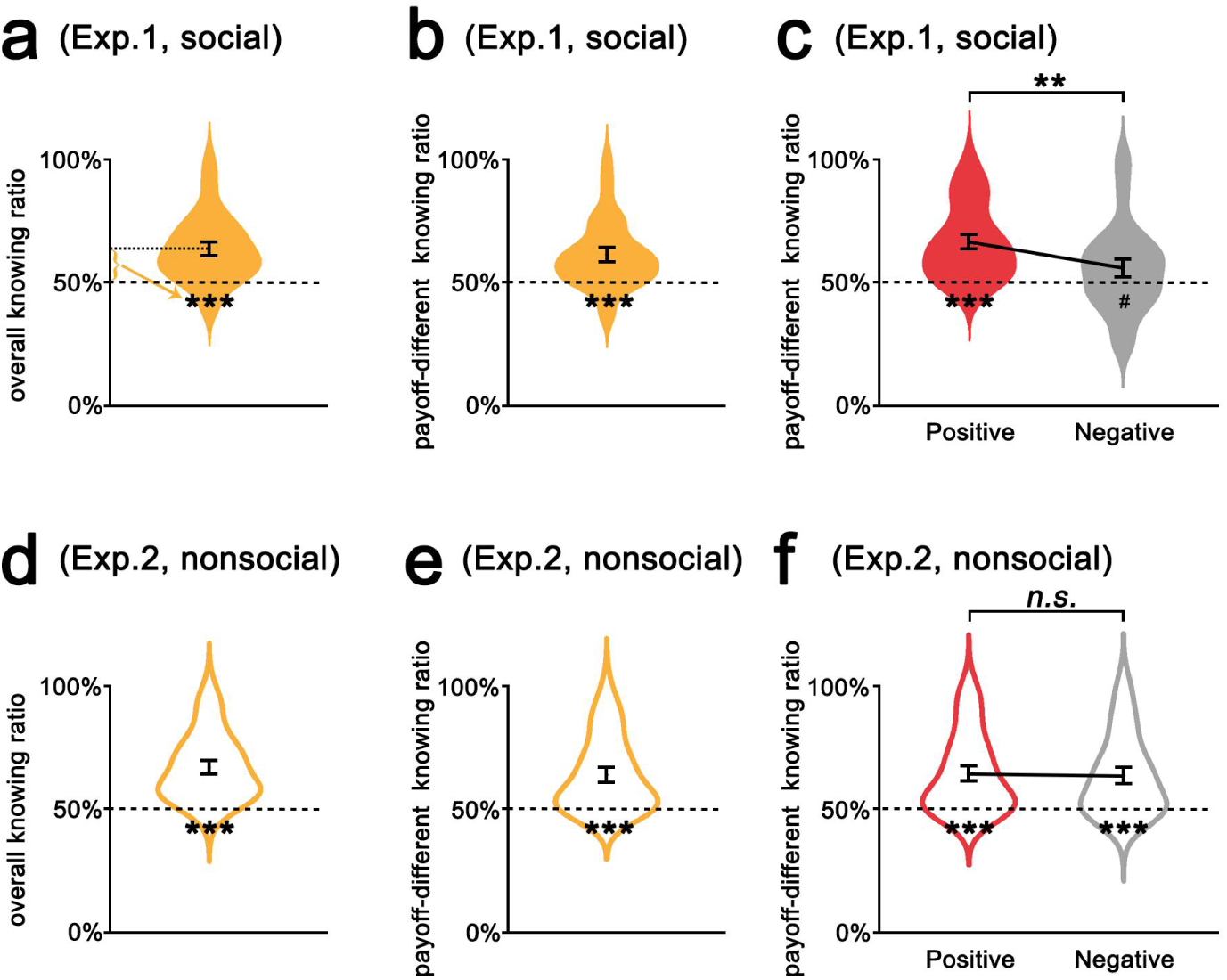
Supplementary model-free results of Exp. 1-2. By calculating the overall and payoff-different knowing ratio, we consistently found that participants preferred to forgo monetary rewards to know both social (***a,b***) and nonsocial evaluations (***d,e***). They preferred to pay more to know social evaluations of positive than negative aspects (66.45 ± 14.87% vs. 55.83 ± 18.14%, *t*(35) = 3.39, *p* = 0.002, ***c***), whereas they would forgo similar amounts of money for the opportunity to know positive and negative non-social evaluations (64.31 ± 15.73% vs. negative: 63.48 ± 15.90%, *t*(34) = 0.55, *p* = 0.584, ***f***). The violin plots indicate the distribution of indices from the pay-to-know task, with elements inside the violin plots representing the mean and standard error. (#*p* < 0.08 and \*\*\**p* < 0.001; *n.s.*, not significant)

**Figure S2.**
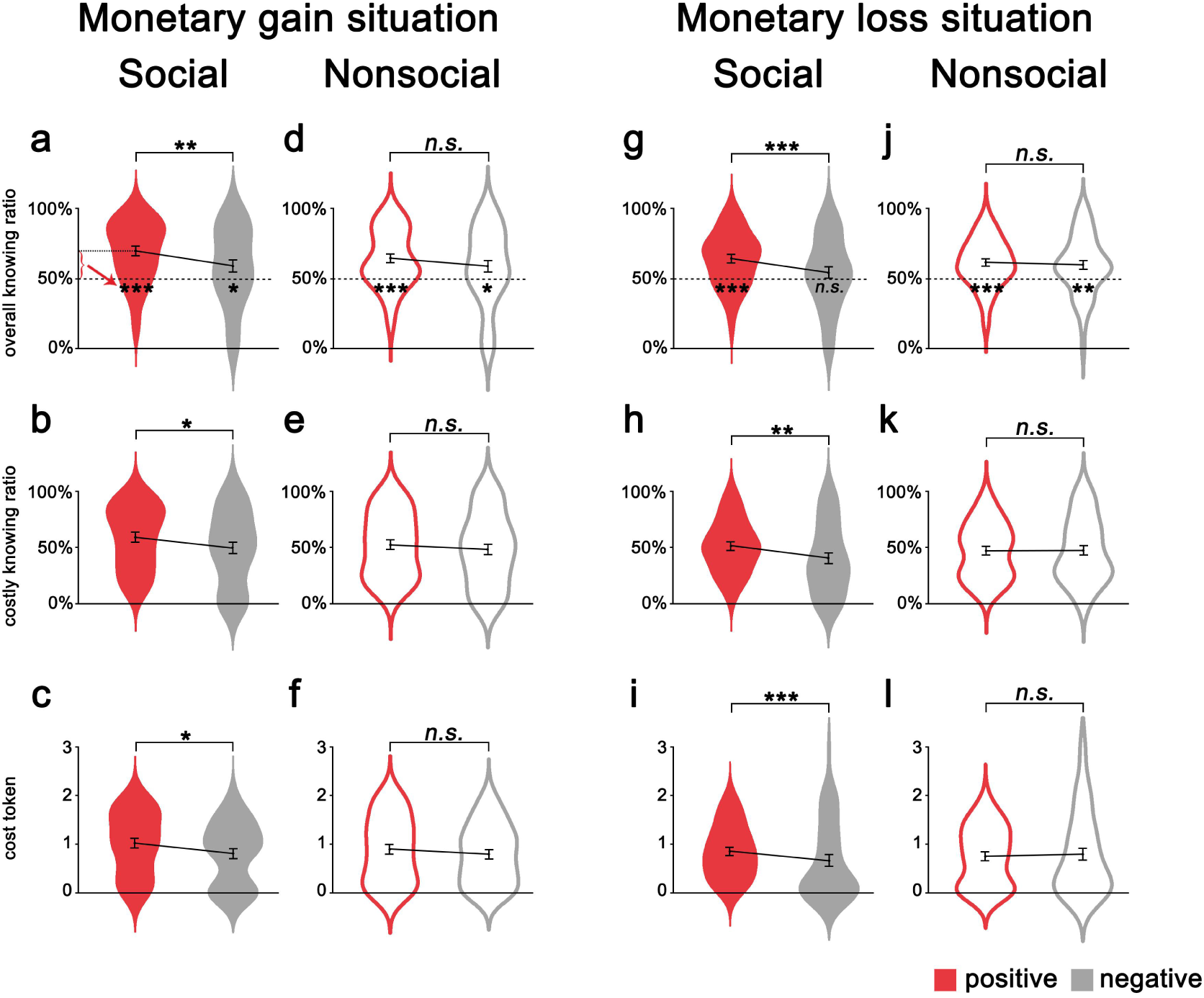
Model-free results for monetary gain and loss situations in Exp. 3. Participants preferred to pay more to know social evaluations of positive aspects than negative aspects in both monetary gain (***a-c***) and loss situations (***g-i***), whereas they would forgo a similar amount of money for the opportunity to know positive and negative non-social evaluations, also in monetary gain (***d-f***) and loss situations (***j-l***). The violin plots indicate the distribution of indices from the pay-to-know task, with elements inside the violin plots representing the mean and standard error. (\**p* < 0.05, \*\**p* < 0.01 and \*\*\**p* < 0.001; *n.s.*, not significant)

**Figure S3.**
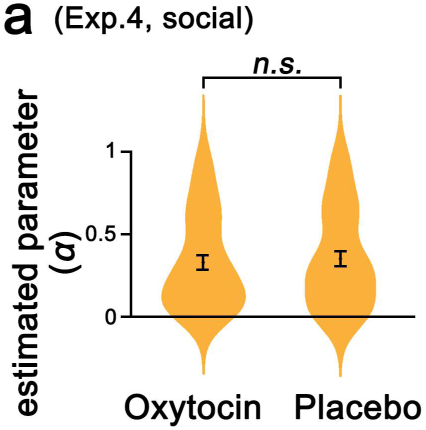
Oxytocin effect on the valuation of social and nonsocial evaluations. No significant effect of oxytocin was found for contribution of monetary payoff difference on the action choice (*α*) when choosing whether to know social evaluation. (*n.s.*, not significant from the bootstrap test).

